# Binding affinity prediction for protein-ligand complex using deep attention mechanism based on intermolecular interactions

**DOI:** 10.1101/2021.03.18.436020

**Authors:** Sangmin Seo, Jonghwan Choi, Sanghyun Park, Jaegyoon Ahn

## Abstract

Accurate prediction of protein-ligand binding affinity is important in that it can lower the overall cost of drug discovery in structure-based drug design. For more accurate prediction, many classical scoring functions and machine learning-based methods have been developed. However, these techniques tend to have limitations, mainly resulting from a lack of sufficient interactions energy terms to describe complex interactions between proteins and ligands. Recent deep-learning techniques show strong potential to solve this problem, but the search for more efficient and appropriate deep-learning architectures and methods to represent protein-ligand complexes continues. In this study, we proposed a deep-neural network for more accurate prediction of protein-ligand complex binding affinity. The proposed model has two important features, descriptor embeddings that contains embedded information about the local structures of a protein-ligand complex and an attention mechanism for highlighting important descriptors to binding affinity prediction. The proposed model showed better performance on most benchmark datasets than existing binding affinity prediction models. Moreover, we confirmed that an attention mechanism was able to capture binding sites in a protein-ligand complex and that it contributed to improvement in predictive performance. Our code is available at https://github.com/Blue1993/BAPA.

**Author summary:** The initial step in drug discovery is to identify drug candidates for a target protein using a scoring function. Existing scoring functions, however, lack the ability to accurately predict the binding affinity of protein-ligand complexes. In this study, we proposed a deep learning-based approach to extract patterns from the local structures of protein-ligand complexes and to highlight the important local structures via an attention mechanism. The proposed model showed good performance for various benchmark datasets compared to existing models.

## Introduction

Structure-based drug design (SBDD) is a method widely used for identification of drug candidates. SBDD includes docking-pose evaluation and estimation of interaction strength between target proteins and small molecules (ligand) [1]. The strength of interactions, known as binding affinity, is generally calculated using various scoring functions. The stronger the interaction, the more the ligand affects the physiological function of the target proteins, so ligands that bind strongly to target protein are selected as drug candidates [2]. Since the predicted binding affinity of the ligand in a library can be used for virtual screening or used to lead optimization, accurate prediction of the binding affinity can reduce the cost of de novo drug design [3].

Mancy scoring functions have been proposed including force-filed-based [4, 5], knowledge-based [6, 7], and empirical scoring functions [8, 9]. Empirical scoring functions are known to show the best prediction performance among the three categories of functions [10], and these empirical functions exploit descriptors of various protein-ligand interactions to calculate a binding affinity score. Those descriptors generally include hydrogen bond, desolvation, van der Waals (vdw), and hydrophobic effects. The coefficient of each descriptor is estimated based on the known binding affinity of protein-ligand complexes. A limitation of the empirical methods, however, is the poor correlation between the experimental and predicted affinity scores. One of the main reasons for this poor correlation is that empirical methods use few terms related to protein-ligand complexes for easy interpretation of results, resulting in failure to describe the true complexity of protein-ligand complexes [11].

Machine learning-based scoring functions [12–16] have been proposed to overcome the limitations of empirical scoring functions and to provide better prediction performance. Those methods exploit various statistical descriptors that are calculated from information about the chemical and physical structure of known protein-ligand complexes [17]. One representative machine learning-based method is RF-Score [12]. This method represents intermolecular interaction by count of atom pairs with nine heavy-atom type (*C*, *N*, *O*, *F*, *P*, *S*, *Cl*, *Br*, and *I*). RF-Score showed significant improvement compared to existing methods on the PDBbind [18] v2007 benchmark set. Moreover, RF-Score v3 [13], which has six additional features, achieved higher predictive accuracy than the original RF-Score model. Another machine learning-based method is structural interaction fingerprints (SIFt) [14], which represents intermolecular interactions with a fingerprint-like format. However, a limitation of SIFt is that the number of interaction types is insufficient to handle the complexity of protein-ligand complexes. To overcome this limitation, structural protein-ligand interaction fingerprints (SPLIF) [15] and protein-ligand extended connectivity (PLEC) fingerprints [16] have been proposed. Both of these methods are based on extended connectivity fingerprints (ECFP) [19].

Recent advances in deep learning in computer vision have led to the development of deep learning-based scoring functions [20–23]. Compared to traditional machine-learning methods, deep learning-based methods do not require domain knowledge for feature selection [24] and can identify hidden patterns using nonlinear transformation [25]. Representative methods using convolutional neural networks (CNN) are Pafnucy [20] and KDEEP [21]. In the two CNN-based methods, each channel is composed of chemical information extracted from a three-dimensional sub-grid for each protein-ligand complex. The problem is that chemical information includes several features such as atomic partial charges, which are calculated using empirical methods like AM1-BCC [26], and which can be incorrect [22].

Fingerprints based on interaction descriptors are an alternative to multidimensional channel representation. However, a limitation of this representation is that it is insufficient to consider the complexity of protein-ligand interaction. We defined descriptors based on RF-Score features for various patterns of interaction. The challenge in this situation, however, is that when diversity is considered, the dimension of fingerprints increases, so it becomes difficult for the predictive model to capture information that is highly related to binding affinity. In sequence-based binding affinity prediction studies, an attention mechanism was introduced to learn binding sites in the training process from training data [27, 28]. We introduced an attention mechanism for capturing descriptors that are important in the prediction of affinity. Another, concern is the lack of distance information in descriptors used in this study. To supplement the descriptors, we used Vina terms, a quantitative numerical value of intermolecular interactions reflecting distance information. This idea was referred to in RF-Score v3.

In the current study, we proposed a deep learning-based model named BAPA (Binding Affinity Prediction with Attention) to improve the accuracy of protein-ligand binding affinity prediction. The proposed model has two important features. First, descriptor embeddings that contains embedded information about the local structures of a protein-ligand complex is learnable, which is means that it is constantly updated for more proper embedding of local structures. Second, we introduced an attention mechanism for highlighting important descriptors to binding affinity prediction. A descriptor vector represents information about the local structure of a protein-ligand complex. In the BAPA, a convolutional layer transforms those descriptor vectors to latent representation and an attention layer captures important descriptors related to binding affinity prediction from the latent representation. This process is shown in Fig 1. When compared with existing methods on four benchmark datasets, BAPA generally show better prediction performance.

**Fig 1.**
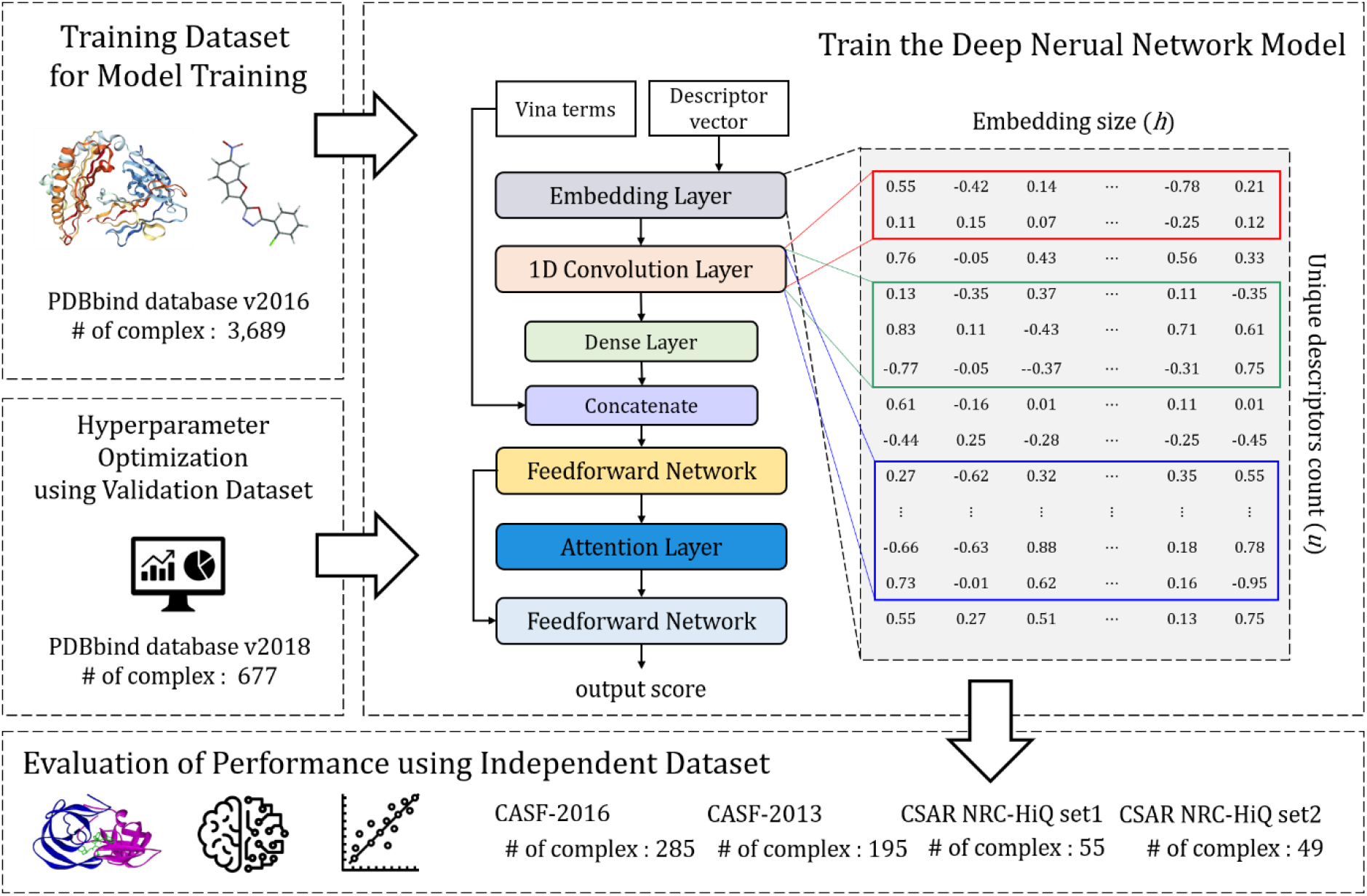
Overview of BAPA experiments. First, we collected training data from the PDBbind database. Second, we constructed a deep learning model that captures local structure patterns that can help predict binding affinity by using convolutional and attention layers. Third, the optimal parameters were found using the validation dataset. Finally, the performance of the model was evaluated using an external test dataset. The numbers (#) of protein-ligand complexes are summarized in each step.

## Results

### Model performance evaluation metrics

The performance of binding affinity prediction models was evaluated via five metrics: mean absolute error (MAE), root mean square error (RMSE), Pearson’s correlation coefficient (*R*_*p*_), Spearman’s correlation coefficient (*R*_*s*_), and standard deviation in regression (SD). MAE and RMSE compute the average of errors between true and predicted affinity scores. Two correlation coefficients measure the linear correlation between true and predicted scores. The SD exhibits the average distance that the true affinities fall from the regression line.

### Selection of the optimal number of descriptors

CNN uses convolution operation to extract features from the input while considering the association of pixels close to the distance (spatial information). However, the order of each element of the descriptor vector *d* is randomly arranged. To arrange the adjacency of related descriptors, we assigned the priority of the descriptors using a random forest regressor trained using the training dataset. Each descriptor is sorted in descending order according to random forest priority.

The initial descriptor vector *d* is a very spare vector, with 9,333 descriptors as elements. The following experiment was conducted to select only the descriptors more essential for the prediction of binding affinity and to represent them in a compact vector form. Among the 9,333 descriptors, the top 500, 1,000, 1,500, 2,000, 2,500 and 3,000 descriptors were selected. After proposed prediction model was trained using the training dataset, and performance evaluation was conducted according to the number of descriptors using the validation dataset. As shown in Table 1, the best performance was shown with the use of 2,500 descriptors, so 2,500 (= *u*) was selected as the optimal number of descriptors.

**Table 1.**
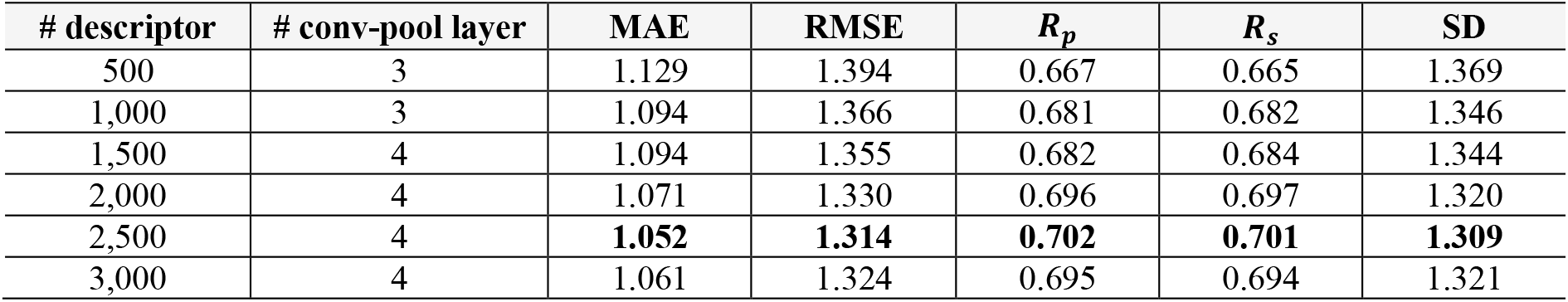
Performance for different number of descriptors.

| # descriptor | # conv-pool layer | MAE | RMSE | $R_p$ | $R_s$ | SD |
| --- | --- | --- | --- | --- | --- | --- |
| 500 | 3 | 1.129 | 1.394 | 0.667 | 0.665 | 1.369 |
| 1,000 | 3 | 1.094 | 1.366 | 0.681 | 0.682 | 1.346 |
| 1,500 | 4 | 1.094 | 1.355 | 0.682 | 0.684 | 1.344 |
| 2,000 | 4 | 1.071 | 1.330 | 0.696 | 0.697 | 1.320 |
| 2,500 | 4 | **1.052** | **1.314** | **0.702** | **0.701** | **1.309** |
| 3,000 | 4 | 1.061 | 1.324 | 0.695 | 0.694 | 1.321 |

### Evaluation of the prediction performance on CASF benchmark

We compared BAPA with four existing popular prediction models including RF-Score v3 [13], Pafnucy [20], PLEC-linear [16], and Onionnet [22]. All models were performance evaluated with the test datasets after training with the training dataset used in this study.

We first presented the results of the CASF-2013 [29] benchmark set containing 195 complexes in Table 2. We confirmed that BAPA outperformed the four-baseline model with *R*_*p*_= 0.77 and *R*_*s*_= 0.77. Furthermore, when compared to the second-best model, BAPA reduced MAE and RMSE by 0.12 and 0.11, respectively.

**Table 2.**
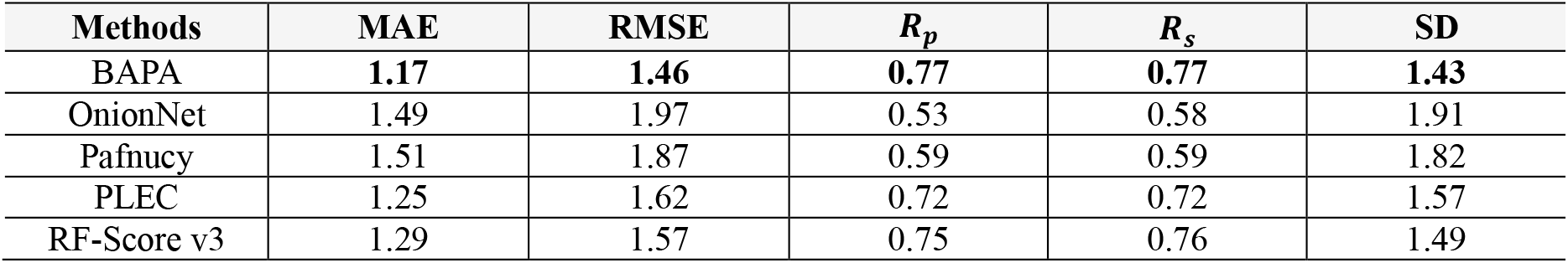
Comparison results using the CASF-2013 dataset.

Results of the CASF-2016 [30] benchmark set containing 285 complexes are shown in Table 3. We confirmed that BAPA has the lowest MAE, RMSE, and SD and highest *R*_*p*_ and *R*_*s*_ compared to other models. Our conclusion for the CASF benchmark sets is that BAPA provided an overwhelming advantage in terms of errors.

**Table 3.** Comparison results using the CASF-2016 dataset.

| Methods | MAE | RMSE | $R_p$ | $R_s$ | SD |
| --- | --- | --- | --- | --- | --- |
| BAPA | **1.02** | **1.31** | **0.82** | **0.82** | **1.25** |
| OnionNet | 1.11 | 1.49 | 0.73 | 0.73 | 1.49 |
| Pafnucy | 1.33 | 1.65 | 0.68 | 0.68 | 1.59 |
| PLEC | 1.14 | 1.45 | 0.76 | 0.75 | 1.41 |
| RF-Score v3 | 1.12 | 1.40 | 0.81 | 0.80 | 1.27 |

### Evaluation of the prediction performance on additional test dataset

We performed additional evaluation of predictive performance using the benchmark set obtained from the external database (CSAR NRC-HiQ [31]). For CSAR NRC-HiQ set2 containing 49 protein-ligand complexes, BAPA showed the best performance with respect to linearity, but with respect to errors, BAPA demonstrated only the second-best performance. These results are shown in Table 4.

**Table 4.** Comparison results using the CSAR NRC-HiQ set2.

| Methods | MAE | RMSE | $R_p$ | $R_s$ | SD |
| --- | --- | --- | --- | --- | --- |
| BAPA | 1.08 | 1.38 | **0.84** | **0.83** | **1.30** |
| OnionNet | 1.55 | 1.97 | 0.62 | 0.69 | 1.86 |
| Pafnucy | 1.52 | 1.91 | 0.68 | 0.66 | 1.74 |
| PLEC | **1.07** | **1.36** | 0.82 | 0.80 | 1.36 |
| RF-Score v3 | 1.26 | 1.61 | 0.78 | 0.76 | 1.47 |

Finally, we presented the results for CSAR NRC-HiQ set1 containing 55 complexes in Table 5. BAPA had the lowest MAE and RMSE and the second-best performance *R*_*p*_, *R*_*s*_, and SD. Our conclusion regarding the CSAR NRC-HiQ set is that it is difficult to say that any specific model is superior, but considering all the metric results, BAPA is slightly superior compared to other models.

**Table 5.** Comparison results using the CSAR NRC-HiQ set1.

| Methods | MAE | RMSE | $R_p$ | $R_s$ | SD |
| --- | --- | --- | --- | --- | --- |
| BAPA | **1.37** | **1.93** | 0.78 | 0.78 | 1.76 |
| OnionNet | 2.29 | 3.03 | 0.35 | 0.47 | 2.64 |
| Pafnucy | 1.83 | 2.43 | 0.59 | 0.55 | 2.29 |
| PLEC | 1.51 | 2.01 | 0.70 | 0.65 | 2.02 |
| RF-Score v3 | 1.45 | 1.99 | **0.80** | **0.86** | **1.70** |

### Generalization performance evaluation based on similarity

In machine learning, generalization refers to the ability of a model to show good performance for new data, that is, data that has not been used for training. In the current study, we attempted to evaluate generalization by constructing complexes of non-overlapping training dataset and test datasets based on PDB ID. However, since the complex is a protein-ligand pair, it should be considered that the proteins of training dataset and homolog proteins or similar proteins may exist in the test datasets. (In the case of a ligand, a homolog ligand or a similar ligand may exist.) Therefore, we performed the following experiment with reference to Li et al., [32] to evaluate the performance of the model according to protein structure similarity or ligand structure similarity.

The structure similarity between the two proteins is calculated by TM-align [33] and is defined as the TM-score [34]. The TM-score has a value between 0 and 1, and higher TM-scores indicate similarity in structure of the two proteins. Since most proteins have multiple chains, the TM-score was measured by comparing all chain structures of each protein. The structure similarity between the two proteins was defined as the lowest TM-score value, since several values may be present depending on the chain structure. Finally, the protein structure similarity of each complex in the test dataset to the training dataset was defined as an average value. Each complex in the test dataset was classified into one of two equal bins, a low similarity bin, and a high similarity bin, according to its similarity with the training dataset.

BAPA showed the best performance regardless of protein structure similarity for three test datasets (CASF-2013, CSAR NRC-HiQ set2, CASF-2016). These results can be seen in Tables 6–7 and S1 Table. In particular, for the high similarity complexes of the CSAR NRC-HiQ set2 test dataset, BAPA showed overwhelmingly high performance compared to other models. The results of the CSAR NRC-HiQ set1 are shown in S2 Table. For the high similarity complexes of the CSAR NRC-HiQ set1 test dataset, RF-Score v3 showed better prediction performance than BAPA. However, BAPA showed better performance than RF-Score v3 for the low structure similarity complexes. Therefore, it may be concluded, that BAPA showed good generalization ability.

**Table 6.** Comparison results using the CASF-2013 with TM-Score.

| Methods | Low Similarity (0.00 – 0.13; 98 complexes) |  |  |  |  | High Similarity (0.13 – 0.18; 97 complexes) |  |  |  |  |
| --- | --- | --- | --- | --- | --- | --- | --- | --- | --- | --- |
| | MAE | RMSE | $R_p$ | $R_s$ | SD | MAE | RMSE | $R_p$ | $R_s$ | SD |
| BAPA | **1.24** | **1.56** | **0.74** | **0.75** | **1.54** | **1.10** | **1.34** | **0.80** | **0.79** | **1.31** |
| OnionNet | 1.46 | 1.91 | 0.59 | 0.64 | 1.85 | 1.51 | 2.03 | 0.46 | 0.51 | 1.94 |
| Pafnucy | 1.57 | 1.94 | 0.59 | 0.59 | 1.84 | 1.43 | 1.78 | 0.58 | 0.59 | 1.78 |
| PLEC | 1.32 | 1.70 | 0.69 | 0.70 | 1.65 | 1.17 | 1.52 | 0.73 | 0.74 | 1.48 |
| RF-Score v3 | 1.35 | 1.65 | 0.73 | 0.74 | 1.57 | 1.23 | 1.48 | 0.77 | 0.77 | 1.38 |

**Table 7.** Comparison results using the CSAR NRC-HiQ set 2 with TM-Score.

| Methods | Low Similarity (0.00 – 0.10; 25 complexes) |  |  |  |  | High Similarity (0.10 – 0.14; 24 complexes) |  |  |  |  |
| --- | --- | --- | --- | --- | --- | --- | --- | --- | --- | --- |
| | MAE | RMSE | $R_p$ | $R_s$ | SD | MAE | RMSE | $R_p$ | $R_s$ | SD |
| BAPA | **0.86** | 1.18 | **0.84** | **0.81** | 1.12 | **1.32** | **1.56** | **0.91** | **0.92** | **0.99** |
| OnionNet | 1.13 | 1.46 | 0.73 | 0.69 | 1.41 | 1.98 | 2.38 | 0.51 | 0.7 | 2.07 |
| Pafnucy | 1.10 | 1.43 | 0.75 | 0.81 | 1.38 | 1.94 | 2.29 | 0.65 | 0.51 | 1.82 |
| PLEC | 1.12 | 1.44 | 0.83 | 0.81 | 1.16 | 1.02 | 1.27 | 0.87 | 0.87 | 1.17 |
| RF-Score v3 | **0.86** | **1.14** | **0.84** | **0.81** | **1.11** | 1.67 | 1.98 | 0.80 | 0.77 | 1.43 |

To evaluate the performance of the model according to the ligand structure similarity, the structure similarity between the two ligands was defined as the SMILES-based Tanimoto coefficient [35]. As with proteins, the ligand structure similarity of each complex of the test dataset to the training dataset was defined as an average value. According to this value, complexes were classified into one of two equal bins of either low similarity or high similarity. In the case of ligands, BAPA showed generally good generalization performance for three test datasets (CASF-2016, CASF-2013, CSAR NRC-HiQ set2). These results are presented in S3-S5 Tables. In the CSAR NRC-HiQ set1 test dataset, BAPA showed lower performance than RF-Score v3 for low similarity complexes (S6 Table). Our conclusion, based on the results of these experiments, is that BAPA shows an overwhelmingly good generalization performance for protein structure similarity but demonstrates limitations in generalization performance for ligand structure similarity. Data on protein structure similarity and ligand structure similarity of each complex used in this experiment are provide in S7-S8 Tables.

### Assessment of module importance via ablation test

BAPA, proposed in this paper, showed good performance in various benchmark sets by applying 1D convolution to input generated from protein-ligand interaction descriptors, adding Vina terms, and applying an attention layer. In this architecture, to analyze the module that most affects the performance of the model, the ablation test was performed assuming 4 cases. The results are shown in Table 8. The descriptor vector is denoted as ‘D’, the attention layer as ‘A’, and Vina terms as ‘V’. In the table, (‘D + A’) means that the experiment was conducted by removing the layer corresponding to the Vina terms from BAPA’s architecture (‘D + V + A’). Similarly, (‘D + V’) means the experiment was performed after removing the attention layer.

**Table 8.** Ablation test results with CASF-2016 dataset.

| Methods | MAE | RMSE | $R_p$ | $R_s$ | SD |
| --- | --- | --- | --- | --- | --- |
| D | 1.15 | 1.45 | 0.77 | 0.77 | 1.39 |
| D + A | 1.12 | 1.41 | 0.78 | 0.78 | 1.36 |
| D + V | 1.04 | 1.34 | 0.80 | 0.80 | 1.29 |
| D + V + A | **1.02** | **1.30** | **0.82** | **0.82** | **1.24** |

The poorest performance was shown when descriptors were used alone, as expected. However, contrary to our expectations, the use of Vina terms (‘D + V’) showed better performance than the use of the attention layer (‘D + A’). In other words, we confirmed that Vina terms have greater influence on predictive performance improvement than the attention layer. However, the best performance was shown when both modules were used, allowing us to confirm that the two modules complement each other and contribute to the performance. Similar trends were shown for the remaining 3 test datasets, as shown in the S9-S11 Tables.

### Analysis of attention vector

BAPA generally showed good performance in test datasets, and we confirmed that the attention layer is an important module for improvement of prediction performance in the ablation test. The presumed explanation for these results was that BAPA identified descriptors that related to the regions of protein-ligand interactions, e.g., binding sites, from the data. To prove this, the attention vectors of the two complexes (PDB ID: 1EBY), (PDB ID: 3DD0) were calculated, and the attention weights corresponding to the top 10% for each complex were then extracted. This has the same meaning as extracting descriptors with the top 10% attention weights for each complex.

The 1EBY complex (HIV protease in complex with the inhibitor BEA369) has 38 binding sites-related interactions based on the sc-PDB database [36]. The inhibitor BEA369 is located in the center and is connected by a purple straight line (ligand bond) in Fig 2A. The green dash lines and the brick-red spoked arcs indicate hydrogen bonds and hydrophobic contacts between the two atoms, respectively. For example, (C48, Asp29-CB) connected by a green dash line indicates a hydrogen bond between the C48 atom of inhibitor BEA369 and the CB atom of aspartic acid, residue 29 of HIV protease. The brick-red spoked arc (C33, Val82-CG1) indicates the hydrophobic contact between the C33 atom of inhibitor BEA369 and the CG1 atom of valine, residue 82 of HIV protease. We confirmed that the extracted top 10% descriptors included all 38 binding sties, and these were highlighted in yellow. These results are presented in Fig 2A. The 3DD0 complex has 9 binding sites-related interactions based on the sc-PDB database. We confirmed that the extracted top 10% descriptors include all interactions except for the two interactions (S2, Val121-CG2) and (N1, Thr199-N). The first refers to the hydrophobic contact between the S2 atom of ethoxzolamide and the CG2 atom of valine, residue 121 of carbonic anhydrase 2, and the second refers to the hydrogen bond between the N1 atom of ethoxzolamide and the N atom of threonine, residue 199 of carbonic anhydrase 2. These results are presented in Fig 2B. We can see that BAPA’s attention layer can capture important regions for interactions. The figure was plotted using Ligplot + [37].

**Fig 2.**
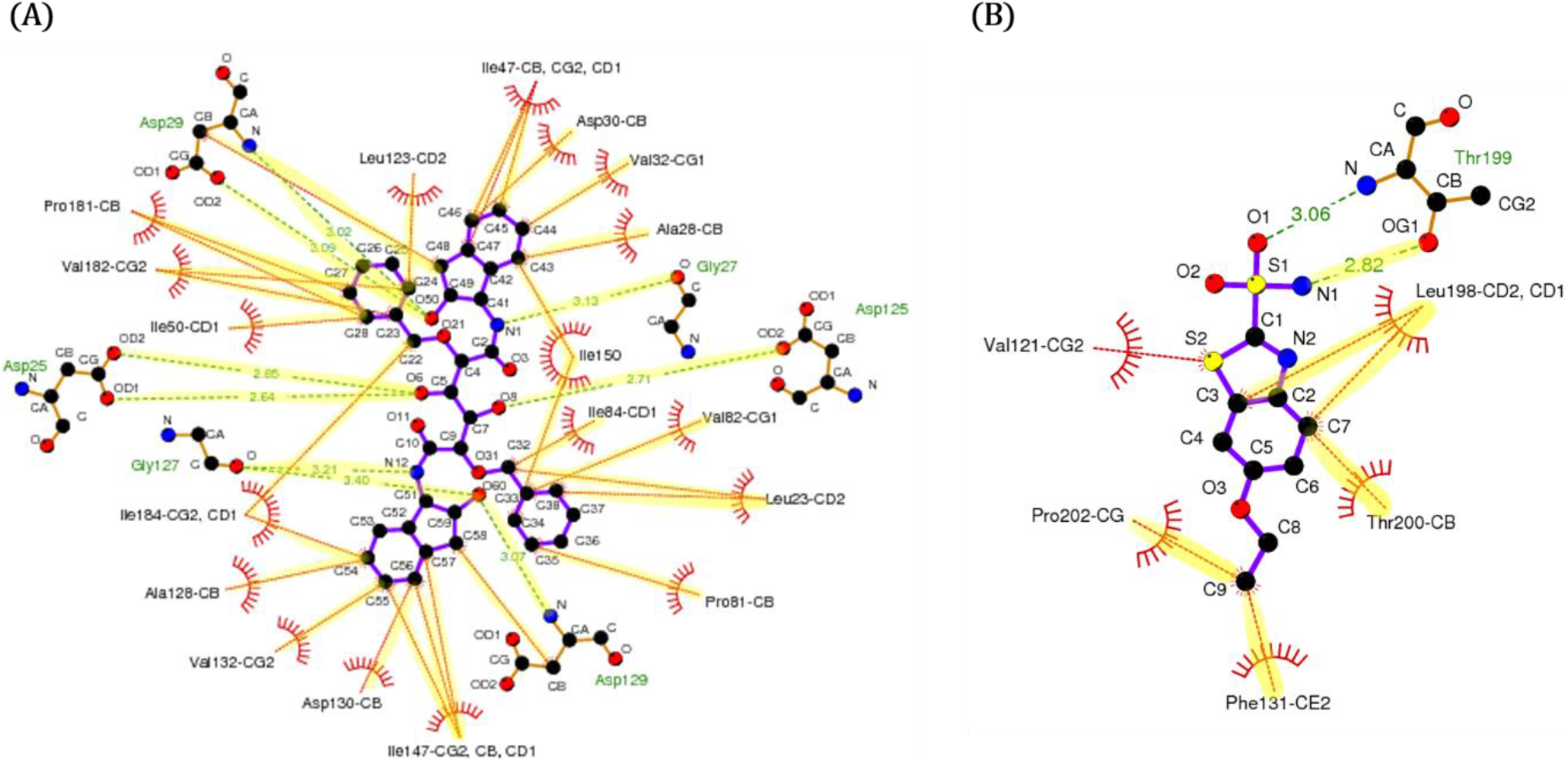
Visualization of interactions sites with high attention score. (A) 1EBY complex, (B) 3DD0 complex. The green dash lines and the brick-red spoked arcs indicate hydrogen bonds and hydrophoic contacts between the two atoms, respectively. Interactions sites captured by BAPA are highlighted in yellow.

## Discussion

In this paper, we proposed BAPA, which could be used for virtual screening and lead optimization in SBDD. The input of convolutional layer was generated using descriptor information and a learnable embedding matrix. The descriptor is a data structure that contains information about the local structure of the protein-ligand complex, and the embedding matrix is a matrix that contains embedded information about the descriptors. The embedding matrix is constantly updated for more appropriate (proper) embedding of the local structure. In addition, the attention mechanism was used to improve the predictive power of the model. The attention model could identify important descriptors in the protein-ligand complex, which would be expected to help researchers design better compounds. BAPA and others existing scoring functions were tested for the CASF-2016 and CASF-2013 benchmark sets. We confirmed that BAPA showed the best performance on both benchmark sets. In addition, BAPA showed good generalization performance for proteins with low structure similarity, which means that more accurate prediction performance can be expected for unseen proteins.

BAPA showed improved predictive performance, but a limitation remains with respect to binding sites that the attention layer cannot capture. There are two main explanations for this limitation. First, we ordered the array of descriptors based on the contribution of random forest in consideration of the convolution operation. However, there may be no association between these sorted descriptors. Second, the architecture of recent neural network models has many layers for feature extraction from the input. For example, Resnet [38] has 15 layers through introduction of skip-connection. BAPA, has a relatively low layer architecture in comparison. In the future, we plan to conduct research to overcome the limitations of our model.

## Materials and Methods

### Building dataset and preprocessing

We collected the PDBbind [18] v2016 *refined set* consisting of 4,507 complexes. The PDBbind dataset consists of 3D structure data of protein-ligand complexes and corresponding binding affinities expressed with *pK*_*a*_ (−*logK*_*d*_ or − *log K*_*i*_) values. We also collected the PDBbind v2018 general set (16,126 complexes), CASF-2016 dataset (285 complexes), and CASF-2013 dataset (195 complexes). The latter two datasets were used only as test datasets for model performance evaluation. Training and validation datasets were constructed using the PDBbind v2016 and v2018 datasets. The training dataset comprised 3,689 complexes obtained by removing complexes that overlap with CASF-2016 and CASF-2013 datasets from the *refined set*. The validation dataset for model parameter optimization was composed of 677 complexes obtained by removing complexes that overlap with the training, CASF-2016, and CASF-2013 datasets from the v2018 *general set*. Removing was processed based on the PDB ID.

Previous studies have suggested that performing training and test using data provided by a specific database tends to yield overly optimized results [39–41]. We collected CSAR NRC-HiQ set1 and CSAR NRC-HiQ set2 composed of 176 and 167 complexes, respectively, as external test datasets. For each dataset, we removed complexes that overlapped with the training, validation, CASF-2016, and CASF-2013 datasets. This resulted in 55 complexes for CSAR NRC-HiQ set1 dataset and 49 complexes for CSAR NRC-HiQ set2 dataset used as test datasets. A summary of each dataset is shown in Fig 1, and PDB IDs for all complexes in each dataset are provided in S12 Table.

In the PDBbind database, the crystal structure data of proteins are provided in PDB file format. All water molecules and cofactors were removed from the crystal structure, and USCF Chimera [42] and Openbabel [43] were used for preprocessing.

### Definition of descriptor

BAPA’s input, a molecular complex, is represented as a 1D vector, which is calculated based on descriptor information obtained from the training dataset. To generate descriptors focused on the contacted protein and ligand atom pairs in the molecular complex, we used nine heavy atoms of the protein and ligand. Let *L* be a list of heavy atoms in ligands [*C*_*L*_, *N_L_*, *O_L_*, *F_L_*, *P_L_*, *S_L_*, *Cl_L_*, *Br_L_*, *I_L_*] where *L*[*i*] is the *i*-th atom type of the ligand (0 ≤ *i* ≤ 8). Likewise, let *P* be a list of heavy atoms in proteins [*C_P_*, *N_P_*, *O_P_*, *F_P_*, *P_P_*, *S_P_*, *Cl_P_*, *Br_P_*, *I_P_*] where *P*[*j*] is the *j*-th atom type of protein (0 ≤ *j* ≤ 8). For each *i* and *j*, a set of contacts *X*[*i*][*j*] is defined by:

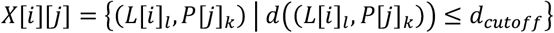

where *L*[*i*]_*l*_ and *P*[*j*]_*k*_ are the *l*-th atom of the *i*-th atom type in the ligand and the *k*-th atom of the *j*-th atom type of the protein, respectively. The distance between the protein atom and the ligand atom pair is calculated by Euclidean distance. We use 12 Å as d_*cutoff*_, based on previous studies [13, 44]. For example, there are two atom pairs with the distance is less than 12 Å (in Fig 3), so *X*[2][2] = {(*L*[2]_4_, *P*[2]_2_)} = {(*O*_*L*4_ ↔ *O*_*P*2_)} and *X*[2][0] = {(*L*[2]_1_, *P*[0]_21_)} = {(*O*_*L*1_ ↔ *C*_*P*21_)}.

**Fig 3.**
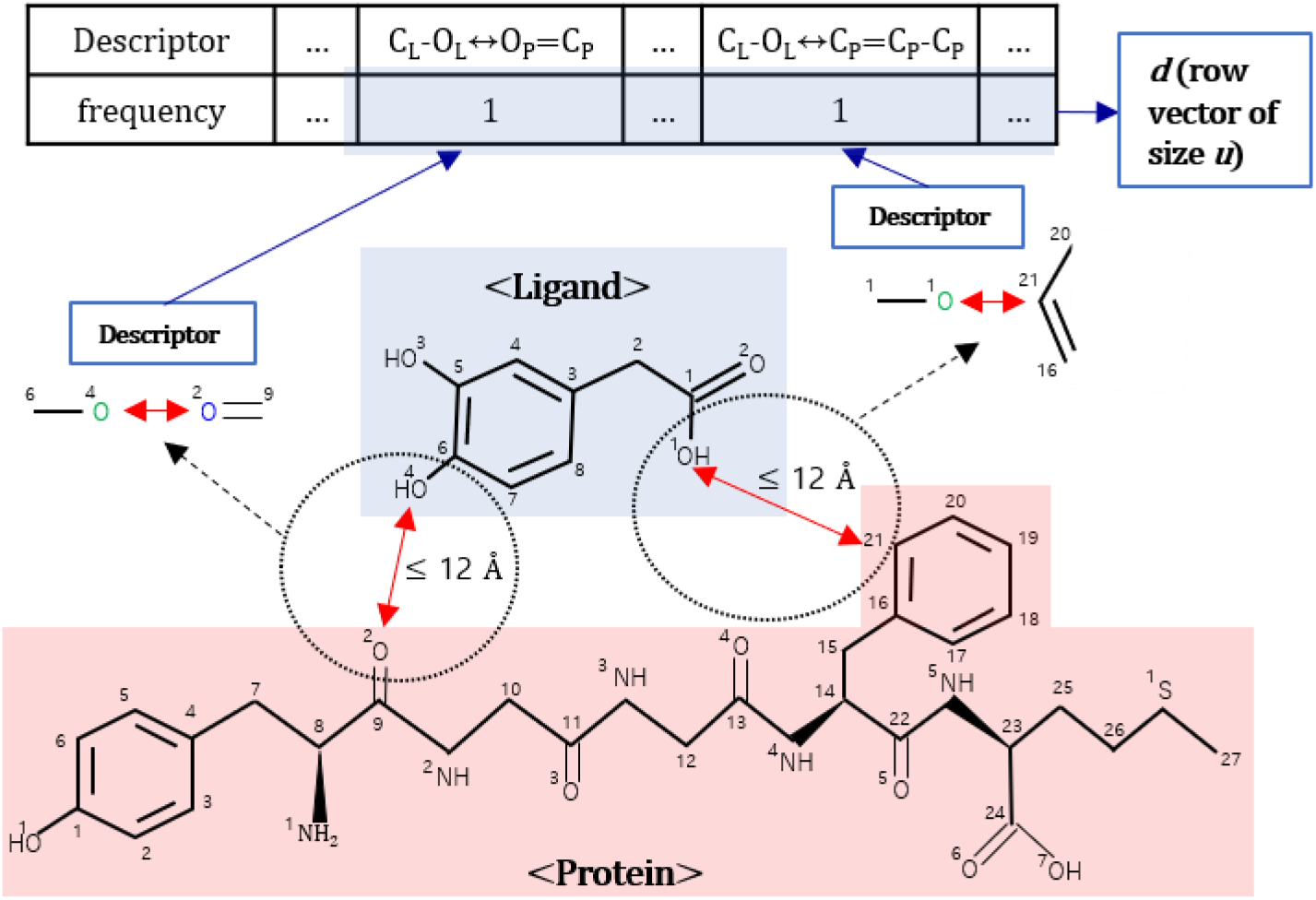
Example of descriptor. There are two atom pairs between which the distance is less than 12 Å, and from which two descriptors can be identified. The frequencies in the protein-ligand complex are counted for all the unique descriptors in the training dataset, with the result being *d*. Note that *OH* is regarded as *O*, and atoms that are not specified are carbon *C*.

The elements of sets *X*[*i*][*j*] form an imaginary edge of which two nodes are ligand and protein atom types. A graph that is extended with one-step neighborhoods from the imaginary edge is defined as a descriptor. The edge between the extended one-step neighborhoods and the imaginary edge has one of five bond types (single, double, triple, amide, and aromatic). Following the previous example, (*O*_*L*4_ ↔ *O*_*P*2_) is extended to ′*C*_*L*6_ − (*O*_*L*4_ ↔ *O*_*P*2_) = *C*_*P*9_′ and (*O*_*L*1_ ↔ *C*_*P*21_) is extended to ′*C*_*L*1_ − (*O*_*L*1_ ↔ *C*_*P*21_) = *C*_*P*20_ − *C*_*P*16_′. Because the order of the bonds (edges) is not considered, ′*C*_*L*1_ − (*O*_*L*1_ ↔ *C*_*P*21_) = *C*_*P*20_ − *C*_*P*16_′ and ′*C*_*L*1_ − (*O*_*L*1_ ↔ *C*_*P*21_) − *C*_*P*16_ = *C*_*P*20_′ are the same. Removal of the atom indexes yields two descriptors, ′*C*_*L*_ − (*O*_*L*_ ↔ *O*_*P*_) = *C*_*P*_′ and ′*C*_*L*_ − (*O*_*L*_ ↔ *C*_*P*_) = *C*_*P*_ − *C*_*P*_′ in Fig 3. In this way, *u* unique descriptors were calculated from the training dataset, and each protein-ligand complex was represented as a descriptor vector *d* having the frequencies of *u* unique descriptors as elements.

### Autodock Vina-based additional features

BAPA exploits Vina terms that reflect distance information of intermolecular interactions in a protein-ligand complex. We used five additional intermolecular Vina terms and one flexible Vina terms. Intermolecular Vina terms consist of three steric interactions (*gauss_1_*, *gauss_2_*, *repulsion*), *hydrophobic*, and *hydrogen bond* terms. The flexible term is the number of active rotatable bonds between the heavy atoms of ligand [45].

### Prediction model

#### Overall schema of deep neural network

The proposed model, BAPA, has three kinds of neural network layers (convolutional, attention, and dense) for binding affinity prediction. We designed the model to extract local structure patterns from descriptor vector *d* via the convolutional layer. The latent representation (encoded vector; *e*) of the complex is calculated from the output of the convolutional layer and Vina terms via feedforward network. Based on this information, the attention layer calculates the descriptors important for affinity prediction and yields an encoded context vector *z*. Finally, the concatenation of an encoded vector *e* and an encoded context vector *z* is input to the feedforward network to predict the binding affinity. Every layer was activated with the exponential linear unit (ELU) function and the whole network was implemented by TensorFlow (1.12.0). The overall architecture is depicted in Fig 4.

**Fig. 4.**
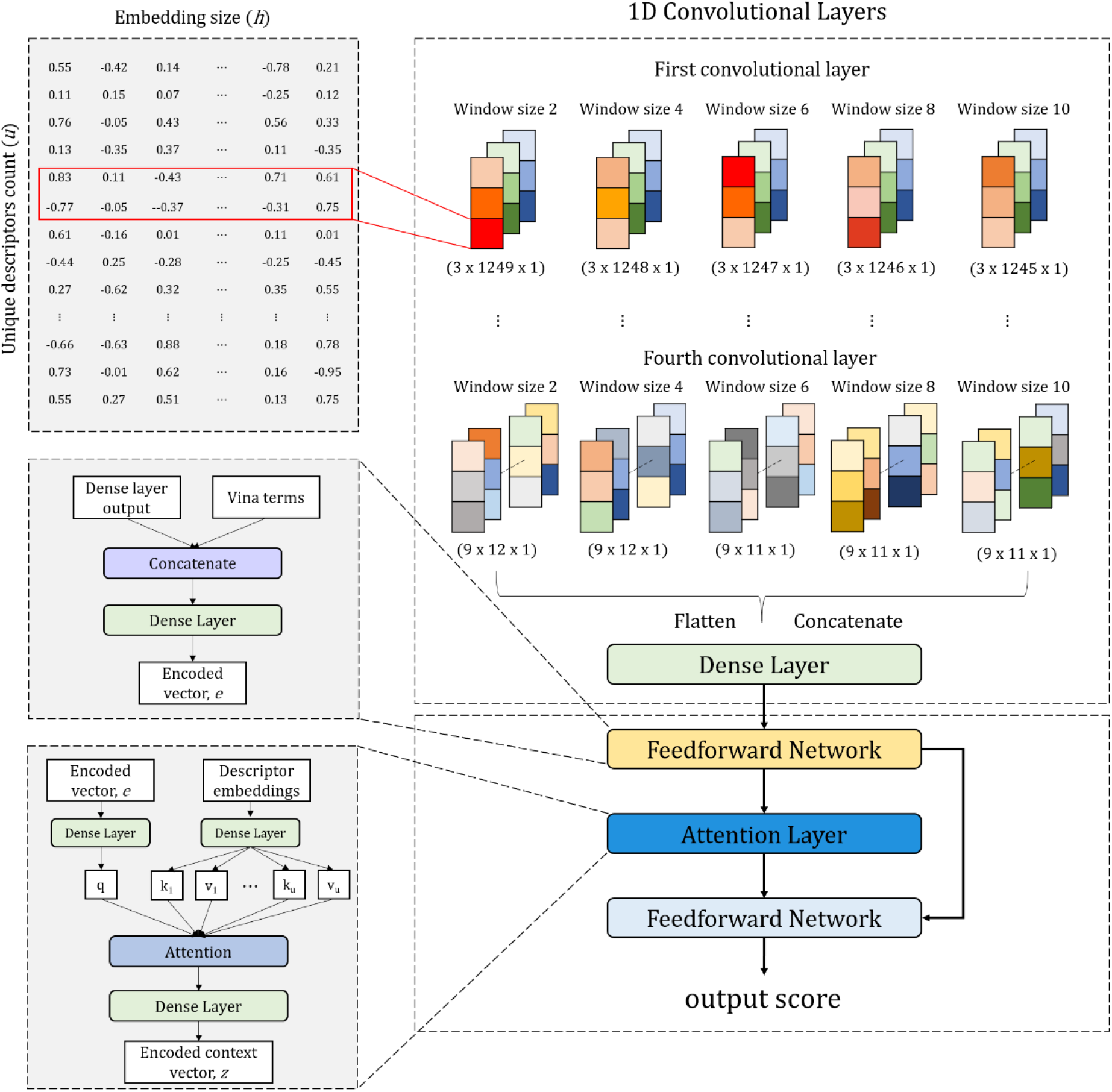
Overview of BAPA. Information about the interaction between the protein and the ligand is represented as a single vector of size 513 via convolutional layer and one dense layer. This vector and the Vina terms are used to generate an encoded vector *e*. The priorities (probabilities) of the descriptors are calculated through the attention layer. Finally, two vectors, the encoded context vector *e,* and the encoded vector *z* are input to the feedforward network and used to predict the affinity.

#### Convolutional layer with descriptor embedding vector

The model starts with the embedding matrix *E* ∈ ℝ^*u* × *h*^ to transform each descriptor to the corresponding embedding vector *E*_*i*_ ∈ ℝ^*h*^ where *u* is the count of descriptors and *h* is the embedding size. An embedding matrix is initialized by the truncated standard normal distribution. To add local structure information to each descriptor embedding vector, an element of the corresponding descriptor vector (frequency of each descriptor) is multiplied by a weight. Then, the input of the convolutional layer is generated through a dense layer for each column of embedding matrix to which local structure information was added.

To find the pattern in the input, all convolutional layers applied one-dimensional (1D) convolution operation. For example, the first convolutional layer uses three filters, and the stride size of each filter is one, so a feature map with a size of (3 × *N* × 1) is generated. To extract various patterns of the descriptors, five different window sizes (2, 4, 6, 8, and 10) were used, so that five (3 × *N* × 1) feature maps were generated in the first convolutional layer. Each of these five feature maps passes through the max pooling layer and decreases in size. The depth of the convolutional layer is four and the convolution operation is the same fashion for all convolutional layers, except that the number of filters is six for the second and third convolutional layers, and nine for the fourth convolutional layer. Detailed parameters for the convolutional layers are provided in S13 Table. The results of the fourth convolutional layer are flattened and concatenated, resulting in a single vector of size 513. The single vector and Vina terms are merged into the encoded vector *e*, which is the latent vector of the complex in the feedforward network.

#### Attention layers for important descriptors

In machine translation, the attention mechanism is mainly designed to solve the problem of long-term dependencies when the input sequence is long. When a word is predicted using a decoder, an attention mechanism puts more focus on words that are more related. In this study, we designed the attention layer to focus on more relevant descriptors. The latent representation of the complex (encoded vector *e*) is input as an attention layer to calculate the contribution of each descriptor to the affinity prediction.

Encoded vector *e* and each row of embedding matrix *E*_*i*_ are calculated into query vector *q*, key vector *k*_i_, and value vector *v*_i_ through a dense layer. Note that in this study the key vector *k*_i_ and the value vector *v*_i_ have the same value. The similarity between query vector *q* and key vector *k*_i_ (0 ≤ *i* ≤ *u*) is calculated using the inner product. The similarities are transformed into descriptor weights via softmax. The weighted sum of the value vector *v*_i_ over the descriptor weight is used as the context vector *c*. The context vector *c* is input to one dense layer to generate the encoded context vector *z*, which is used to predict the binding affinity together with encoded vector *e*.

#### Feedforward network for binding affinity

The encoded context vector *z*, which is an output of the attention layer, and the encoded vector *e*, are used to predict the binding affinity (*pK*_*a*_) through the feedforward network consisting of 512, 256, and 128 neurons.

#### Definition of loss function and weight optimization

In the proposed neural network model, the input flows to the output layer in a feedforward fashion. The mean squared error was used as a loss function to train the weights and biases. To prevent overfitting, we applied L2 regularization, so the norm of weights is added to the loss. The Adam optimizer was used for training the network (learning rate 0.005, batch size 256).

## Supporting information

**S1 Table. Comparison results using the CASF-2016 dataset with TM-Score.**

**S2 Table. Comparison results using the CSAR-HiQ set1 dataset with TM-Score.**

**S3 Table. Comparison results using the CASF-2013 dataset with Tanimoto coefficient.**

**S4 Table. Comparison results using the CASF-2016 dataset with Tanimoto coefficient.**

**S5 Table. Comparison results using CSAR-HiQ set2 dataset with Tanimoto coefficient.**

**S6 Table. Comparison results using CSAR-HiQ set1 dataset with Tanimoto coefficient.**

**S7 Table. Protein structure similarity of the test dataset to the training dataset with TM-Score.**

**S8 Table. Ligand structure similarity of the test dataset to the training dataset with Tanimoto coefficient.**

**S9 Table. Ablation test results with CASF-2013 dataset.**

**S10 Table. Ablation test results with CSAR NRC-HiQ set2.**

**S11 Table. Ablation test results with CSAR NRC-HiQ set1.**

**S12 Table. PDB IDs list for all complexes in each dataset.**

**S13 Table. Parameters and architecture in convolutional layers.**

## Author Contributions

**Conceptualization**: Sangmin Seo, Jaegyoon Ahn

**Data curation**: Sangmin Seo

**Formal analysis**: Sangmin Seo

**Funding acquisition:** Sanghyun Park, Jaegyoon Ahn

**Investigation**: Sangmin Seo

**Methodology**: Sangmin Seo, Jaegyoon Ahn

**Project administration**: Sanghyun Park, Jaegyoon Ahn

**Resources**: Sanghyun Park, Jaegyoon Ahn

**Software**: Sangmin Seo

**Supervision**: Sanghyun Park, Jaegyoon Ahn

**Validation**: Sangmin Seo, Jonghwan Choi

**Visualization**: Sangmin Seo, Jonghwan Choi

**Writing – original draft**: Sangmin Seo, Jonghwan Choi, Sanghyun Park, Jaegyoon Ahn

**Writing – review & editing**: Sangmin Seo, Jonghwan Choi, Sanghyun Park, Jaegyoon Ahn

## References

1. Kroemer RT. Structure-based drug design: docking and scoring. Current protein and peptide science. 2007;8(4):312–28.

2. Li S, Xi L, Wang C, Li J, Lei B, Liu H, et al. A novel method for protein-ligand binding affinity prediction and the related descriptors exploration. Journal of computational chemistry. 2009;30(6):900–9.

3. DiMasi JA, Hansen RW, Grabowski HG. The price of innovation: new estimates of drug development costs. Journal of health economics. 2003;22(2):151–85.

4. Ewing TJ, Makino S, Skillman AG, Kuntz ID. DOCK 4.0: search strategies for automated molecular docking of flexible molecule databases. Journal of computer-aided molecular design. 2001;15(5):411–28.

5. Jones G, Willett P, Glen RC, Leach AR, Taylor R. Development and validation of a genetic algorithm for flexible docking. Journal of molecular biology. 1997;267(3):727–48.

6. Muegge I. PMF scoring revisited. Journal of medicinal chemistry. 2006;49(20):5895–902.

7. Velec HF, Gohlke H, Klebe G. DrugScoreCSD knowledge-based scoring function derived from small molecule crystal data with superior recognition rate of near-native ligand poses and better affinity prediction. Journal of medicinal chemistry. 2005;48(20):6296–303.

8. Gehlhaar DK, Verkhivker GM, Rejto PA, Sherman CJ, Fogel DR, Fogel LJ, et al. Molecular recognition of the inhibitor AG-1343 by HIV-1 protease: conformationally flexible docking by evolutionary programming. Chemistry & biology. 1995;2(5):317–24.

9. Wang R, Lai L, Wang S. Further development and validation of empirical scoring functions for structure-based binding affinity prediction. Journal of computer-aided molecular design. 2002;16(1):11–26.

10. Cheng T, Li X, Li Y, Liu Z, Wang R. Comparative assessment of scoring functions on a diverse test set. Journal of chemical information and modeling. 2009;49(4):1079–93.

11. Li G-B, Yang L-L, Wang W-J, Li L-L, Yang S-Y. ID-Score: a new empirical scoring function based on a comprehensive set of descriptors related to protein–ligand interactions. Journal of chemical information and modeling. 2013;53(3):592–600.

12. Ballester PJ, Mitchell JB. A machine learning approach to predicting protein–ligand binding affinity with applications to molecular docking. Bioinformatics. 2010;26(9):1169–75.

13. Li H, Leung KS, Wong MH, Ballester PJ. Improving AutoDock Vina using random forest: the growing accuracy of binding affinity prediction by the effective exploitation of larger data sets. Molecular informatics. 2015;34(2-3):115–26.

14. Deng Z, Chuaqui C, Singh J. Structural interaction fingerprint (SIFt): a novel method for analyzing three-dimensional protein− ligand binding interactions. Journal of medicinal chemistry. 2004;47(2):337–44.

15. Da C, Kireev D. Structural protein–ligand interaction fingerprints (SPLIF) for structure-based virtual screening: method and benchmark study. Journal of chemical information and modeling. 2014;54(9):2555–61.

16. Wójcikowski M, Kukiełka M, Stepniewska-Dziubinska MM, Siedlecki P. Development of a protein– ligand extended connectivity (PLEC) fingerprint and its application for binding affinity predictions. Bioinformatics. 2019;35(8):1334–41.

17. Manly CJ, Louise-May S, Hammer JD. The impact of informatics and computational chemistry on synthesis and screening. Drug discovery today. 2001;6(21):1101–10.

18. Wang R, Fang X, Lu Y, Wang S. The PDBbind database: Collection of binding affinities for protein− ligand complexes with known three-dimensional structures. Journal of medicinal chemistry. 2004;47(12):2977–80.

19. Rogers D, Hahn M. Extended-connectivity fingerprints. Journal of chemical information and modeling. 2010;50(5):742–54.

20. Stepniewska-Dziubinska MM, Zielenkiewicz P, Siedlecki P. Development and evaluation of a deep learning model for protein–ligand binding affinity prediction. Bioinformatics. 2018;34(21):3666–74.

21. Jiménez J, Skalic M, Martinez-Rosell G, De Fabritiis G. K deep: protein–ligand absolute binding affinity prediction via 3d-convolutional neural networks. Journal of chemical information and modeling. 2018;58(2):287–96.

22. Zheng L, Fan J, Mu Y. Onionnet: a multiple-layer intermolecular-contact-based convolutional neural network for protein–ligand binding affinity prediction. ACS omega. 2019;4(14):15956–65.

23. Nguyen DD, Wei G-W. AGL-Score: Algebraic graph learning score for protein–ligand binding scoring, ranking, docking, and screening. Journal of chemical information and modeling. 2019;59(7):3291–304.

24. Schmidhuber J. Deep learning in neural networks: An overview. Neural networks. 2015;61:85–117.

25. LeCun Y, Bengio Y, Hinton G. Deep learning. nature. 2015;521(7553):436–44.

26. Jakalian A, Jack DB, Bayly CI. Fast, efficient generation of high-quality atomic charges. AM1-BCC model: II. Parameterization and validation. Journal of computational chemistry. 2002;23(16):1623–41.

27. Zhao Q, Xiao F, Yang M, Li Y, Wang J, editors. AttentionDTA: prediction of drug–target binding affinity using attention model. 2019 IEEE International Conference on Bioinformatics and Biomedicine (BIBM); 2019: IEEE.

28. Karimi M, Wu D, Wang Z, Shen Y. DeepAffinity: interpretable deep learning of compound–protein affinity through unified recurrent and convolutional neural networks. Bioinformatics. 2019;35(18):3329–38.

29. Li Y, Liu Z, Li J, Han L, Liu J, Zhao Z, et al. Comparative assessment of scoring functions on an updated benchmark: 1. Compilation of the test set. Journal of chemical information and modeling. 2014;54(6):1700–16.

30. Su M, Yang Q, Du Y, Feng G, Liu Z, Li Y, et al. Comparative assessment of scoring functions: the CASF-2016 update. Journal of chemical information and modeling. 2018;59(2):895–913.

31. Dunbar Jr JB, Smith RD, Damm-Ganamet KL, Ahmed A, Esposito EX, Delproposto J, et al. CSAR data set release 2012: ligands, affinities, complexes, and docking decoys. Journal of chemical information and modeling. 2013;53(8):1842–52.

32. Li Y, Yang J. Structural and sequence similarity makes a significant impact on machine-learning-based scoring functions for protein–ligand interactions. Journal of chemical information and modeling. 2017;57(4):1007–12.

33. Zhang Y, Skolnick J. TM-align: a protein structure alignment algorithm based on the TM-score. Nucleic acids research. 2005;33(7):2302–9.

34. Zhang Y, Skolnick J. Scoring function for automated assessment of protein structure template quality. Proteins: Structure, Function, and Bioinformatics. 2004;57(4):702–10.

35. Rácz A, Bajusz D, Héberger K. Life beyond the Tanimoto coefficient: similarity measures for interaction fingerprints. Journal of cheminformatics. 2018;10(1):1–12.

36. Desaphy J, Bret G, Rognan D, Kellenberger E. sc-PDB: a 3D-database of ligandable binding sites—10 years on. Nucleic acids research. 2015;43(D1):D399–D404.

37. Wallace AC, Laskowski RA, Thornton JM. LIGPLOT: a program to generate schematic diagrams of protein-ligand interactions. Protein engineering, design and selection. 1995;8(2):127–34.

38. Targ S, Almeida D, Lyman K. Resnet in resnet: Generalizing residual architectures. arXiv preprint arXiv:160308029. 2016.

39. Gabel J, Desaphy J, Rognan D. Beware of Machine Learning-Based Scoring Functions On the Danger of Developing Black Boxes. Journal of chemical information and modeling. 2014;54(10):2807–15.

40. Kramer C, Gedeck P. Leave-cluster-out cross-validation is appropriate for scoring functions derived from diverse protein data sets. Journal of chemical information and modeling. 2010;50(11):1961–9.

41. Ballester PJ, Mitchell JB. Comments on “leave-cluster-out cross-validation is appropriate for scoring functions derived from diverse protein data sets”: Significance for the validation of scoring functions. ACS Publications; 2011.

42. Pettersen EF, Goddard TD, Huang CC, Couch GS, Greenblatt DM, Meng EC, et al. UCSF Chimera—a visualization system for exploratory research and analysis. Journal of computational chemistry. 2004;25(13):1605–12.

43. O’Boyle NM, Banck M, James CA, Morley C, Vandermeersch T, Hutchison GR. Open Babel: An open chemical toolbox. Journal of cheminformatics. 2011;3(1):1–14.

44. Ballester PJ, Schreyer A, Blundell TL. Does a more precise chemical description of protein–ligand complexes lead to more accurate prediction of binding affinity? Journal of chemical information and modeling. 2014;54(3):944–55.

45. Trott O, Olson AJ. AutoDock Vina: improving the speed and accuracy of docking with a new scoring function, efficient optimization, and multithreading. Journal of computational chemistry. 2010;31(2):455–61.

